# Multiple variants of IncF plasmid alleles discovered within single bacterial cells challenge previous assumptions

**DOI:** 10.1101/2024.07.26.605309

**Authors:** Michaela Ruzickova, Jana Palkovicova, Ivo Papousek, Max L Cummins, Steven P Djordjevic, Monika Dolejska

**Affiliations:** Central European Institute of Technology, University of Veterinary Sciences Brno, Brno, Czech Republic; Department of Biology and Wildlife Diseases, University of Veterinary Sciences Brno, Brno, Czech Republic; Biomedical Centre, Charles University, Pilsen, Czech Republic; Australian Institute for Microbiology and Infection, University of Technology Sydney, Australia; The Australian Centre for Genomic Epidemiological Microbiology, University of Technology Sydney, Australia; Department of Clinical Microbiology and Immunology, Institute of Laboratory Medicine, The University Hospital Brno, Czech Republic

## Abstract

IncF plasmids are diverse mobile genetic elements found in bacteria from the Enterobacteriaceae family and often carry critical antibiotic and virulence gene cargo. The classification of IncF plasmids using the plasmid Multi-Locus Sequence Typing (pMLST) tool compares the sequences of IncF alleles against a database to create a plasmid sequence type (ST). Accurate identification of plasmid STs is epidemiologically useful because it enables an assessment of the crucial IncF plasmid lineages associated with pandemic and emerging enterobacterial sequence types, inferring important information about specific bacterial lineages. Our initial observations showed discrepancies in IncF allele variants reported by pMLST in a collection of 898 *Escherichia coli* ST131 genomes. To evaluate the limitations of the pMLST tool, we interrogated an in-house and publicly available repository of 70324 *E. coli* genomes of various STs and other Enterobacterales genomes (n=1247). All short-read genomes and representatives selected for long-read sequencing were used to assess allele variants and to compare the output with the real biological situation. When multiple allele variants occurred in a single bacterial genome, the python and web versions of the tool randomly selected one allele to report, leading to limited and inaccurate ST identification. Discrepancies were detected in 5804 of 72469 genomes (8.01%). Long read sequencing of 27 carefully selected genomes confirmed multiple IncF allele variants present on one plasmid, or two separate IncF plasmids present in a single bacterial cell. The pMLST tool was unable to accurately distinguish allele variants and their location on replicons using short-read nor long-read genome sequences.

**Importance:** Plasmid sequence type is crucial for describing IncF plasmids due to their capacity to carry important antibiotic and virulence gene cargo and consequently due to their association with disease-causing enterobacterial lineages exhibiting resistance to clinically relevant antibiotics in humans and food animals. As a result, precise reporting of IncF allele variants in IncF plasmids is necessary. Comparison of the FAB formulae generated by the plasmid Multi-Locus Sequence Typing (pMLST) tool with annotated long-read genome sequences identified inconsistencies, including examples where multiple IncF allele variants were present on the same plasmid but missing in the FAB formula, or in cases where two IncF plasmids were detected in one bacterial cell and the pMLST output provided information only about one plasmid. Such inconsistencies may cloud interpretation of IncF plasmid replicon type in specific bacterial lineages or inaccurate assumptions of host strain clonality.

## Introduction

Plasmids of incompatibility group F (IncF) are diverse mobile genetic elements widely found in various bacteria from the Enterobacteriaceae family [1]. Due to their conjugative abilities, they are horizontally transferred among such species, which increases their potential impact on bacterial evolution [2]. IncF plasmids are multi-replicon plasmids that usually carry more than one replicon responsible for the initiation of plasmid DNA replication. The mosaic structure of the IncF plasmids may have implications for broader host range replication [3], [4]. Replicons located on typical IncF plasmids belong to FII, FIA and FIB families with the latter two being the ones predominantly used for the replication initiation, which leaves the FII replicon free to be used when incompatibility obstacles arise. If a cell contains two plasmids, both carrying FII, FIA and FIB replicons, each plasmid can use a different replicon for its replication allowing both of them to be stably maintained in the cell [3], [5].

The Plasmid Multi-Locus Sequence Typing (pMLST) *in silico* tool is often used to further classify plasmids. It targets specific sequences linked to the regions of replication proteins in plasmid genomes and uses BLASTn to compare them against a database of known replicon alleles [6]. To classify IncF plasmids, a scheme called replicon sequence typing (RST) was established within the pMLST tool, based on the variety of known IncF replicon alleles. The output of this analysis, known as the FAB formula, is created by the allele variants for each of the plasmid replicons FII, FIA and FIB, respectively, and can be further used for plasmid identification. However, variations of these three basic replicons also exist within the formula such as in the FII replicon, which belongs to a larger FII family together with FII_K_, FII_S_ and FII_Y_ alleles being mostly species-specific for *Klebsiella* sp., *Salmonella* sp. and *Yersinia* sp., respectively. Its place in the formula can be also taken by the FIC replicon from the same FII family which occurs regardless of the species [5], [7].

Precise calling of replicons and allele variants present in mosaic IncF plasmids is of high importance not only for deciphering their evolution, but also for epidemiology of the clinically relevant plasmid types responsible for dissemination of antibiotic resistance and virulence genes [8]. The FAB formula is a significant parameter for determining the association of IncF plasmids with certain bacterial species and genotypes, and omission of some of the alleles may lead to improper evaluation of clonality among bacterial strains [5], [9].

During the analysis of IncF plasmid content in a large collection of *E. coli* sequence type (ST) 131 isolates, inconsistent results of repeated RST analysis were observed in regard to allele variants of plasmid replicons that define the FAB formula. The main goal of this project was to: i) assess the observed discrepancies; ii) identify affected plasmid lineages; iii) identify specific alleles by analysing an extensive collection of *E. coli* genomes (n=70324) as well as other species from the Enterobacteriaceae family (n=1247) and iv) examine complete plasmid sequences from our in-house collection as well as publicly available databases (n=7441) for inconsistency in the FAB formulae.

## Methodology

### Collection of bacterial genomes

Our initial observations showed the discrepancies in the identified replicon alleles in the in-house collection of Czech *E. coli* ST131 genomes (n=898) from various human (79.40%; 713/898), animal (5.12%; 46/898), and environmental sources (15.48%; 139/898) (partially published under BioProject PRJNA983524 and PRJNA595483 [10]). Since a detailed pMLST analysis pointed out inaccurate report of the IncF allele variants, more sequences were assessed to evaluate aberrant pMLST calling within larger dataset. This was done to exclude the possibility of a technical error and avoid potential bias from our collection originating from a single country and a single *E. coli* ST. These analyses included the international collection of *E. coli* ST131 genomes (n=13656) which was accessed from Enterobase (https://enterobase.warwick.ac.uk/) on 21.2.2023. Moreover, 121427 genomes of various *E. coli* STs were also obtained from Enterobase on 30.5.2023 together with their metadata in order to search for the phenomenon in other *E. coli* STs as well. Sequences with missing metadata (source, country, year) and those which did not pass quality check were excluded from this selection, together with *E. coli* ST131 genomes in case of the various STs collection. Quality checks considered the number of contigs (threshold 400 contigs) and the total length of the genome (threshold 4700000 base pairs). In addition, non-ST131 *E. coli* isolates (n=1743) from our in-house collection were also included. The entire collection sourced from Enterobase comprised of 8514 *E. coli* ST131 genomes and 60067 sequences of other *E. coli* STs.

Lastly, to check for this phenomenon in genera other than *E. coli*, an in-house collection of whole-genome sequences of Enterobacterales (n=2342) accumulated during previous projects was used for the assessment of aberrant IncF plasmid replicon calls. Genomes with unreliable species affiliation and poor quality were excluded, and the final collection consisted of 1247 genomes.

Raw data from short-read sequencing of each in-house collection were quality (Q≥20) and adaptor trimmed and *de novo* assembled which was followed by the quality and contamination check. Downloaded publicly available short-read sequences went through quality and contamination check. The presence of plasmids in these genomes was determined using ABRicate (https://github.com/tseemann/abricate) and database PlasmidFinder with coverage and identity threshold 90% for the presence of plasmid replicons [6].

### Collection of complete plasmid sequences

Complete IncF plasmids (n=7441) were accessed on 27.1.2023 to support our findings. Reference sequences of FII, FIA, FIB, FIC, FII_K_, FII_S_ and FII_Y_ alleles downloaded from PubMLST (https://pubmlst.org/) were used as query sequences to search for the corresponding plasmid genomes in the NCBI database [11]. No filtering based on metadata was done in this collection since we did not intend to use the data for epidemiological comparison. These sequences obtained from a public database were not run through quality check as they were considered complete plasmid sequences. Data from all of the collections can be seen in the supplement table S1.

To confirm presumptions based on short-read sequence analyses, accurately determine plasmid replicons, and to obtain complete plasmid sequences, several representative *E. coli* ST131 isolates from the in-house collection (n=27) were also subjected to long-read sequencing. The representatives were selected based on the multiple allele variants detected in the short-read genome assemblies, presence of antibiotic resistance genes, and their clonality. This was necessary to confirm the presence of alleles identified by pMLST and identify the location of multiple allele variants in the genome. Genomic DNA of the selected isolates was extracted and used for the preparation of sequence libraries according to the manufacturer’s protocol. The mixture was subsequently loaded onto a flow cell and sequenced on a MinION platform (Oxford Nanopore Technologies, UK). The long-read raw data obtained by base-calling under high-accuracy model were adaptor-trimmed, demultiplexed and quality-trimmed (Q≥9). The long reads were *de novo* assembled and obtained assemblies were polished using both corrected long reads and corrected short reads. Obtained genomes were used for the detection of IncF allele variants. The plasmids then underwent automatic annotation with manual supervision. Annotated long-read sequences are available on GenBank under accession numbers PQ066878-PQ066914. All the kits, platforms and software used for both short and long-read sequencing and data processing are described in the supplement table S1.

### pMLST tool versions used for the analyses

All genome and plasmid sequences were subjected to IncF plasmid typing by pMLST analysis using three different versions of the pMLST tool. Web-based tool pMLST 2.0 from the Center for Genomic Epidemiology (https://www.genomicepidemiology.org/) as well as two versions of downloadable python pMLST software (https://bitbucket.org/genomicepidemiology/pmlst/) were used for the RST of IncF plasmids. The web-based tool provides a table of hits for each allele including identity and coverage values and creates the FAB formula from the hits. The two python versions were pulled either from Docker (https://github.com/docker) or Anaconda (https://github.com/conda) repositories. Output of both versions are text files for each of the analysed isolates which show the allele hits together with their identity and coverage and suggest the FAB formula. The FAB formula is then reported as a “Sequence type” of the plasmid by all of the pMLST tools.

## Results and discussion

### Various pMLST tools are not able to sufficiently distinguish multiple IncF replicons

Inconsistent results regarding the FAB formula were observed during repeated pMLST analysis using three different versions of pMLST tool. All of them are capable of finding two of the allele variants if they are both present in either one of the FII, FIA and FIB replicons. In case that more than one allele occurs in a single replicon, a warning message “FII/FIA/FIB: Multiple perfect hits found” appears beneath the output. However, a single resulting FAB formula presented by all versions of the tool as an ST of the analysed plasmid is created with a random selection of one of such multiple hits and thus does not reflect the allele content found by the tool and the accurate prediction of specific plasmid(s). Instances of genomes carrying three different allele variants or two copies of the same allele variant on a single FII, FIA or FIB replicon were also observed, however, these were not confirmed by long-read sequencing. These situations were both detected only by the web-based version of the tool while surprisingly, the python versions did not display the extra alleles. The information about the multiple FII allele hits is stored in the detailed results of the pMLST analysis in all three versions of the tool, see supplement files S2-S4. This information can be extracted and added manually to an FAB formula but is not automatically included in the final plasmid ST. Even though we extract it manually, this information does not necessary enable an accurate identification of the IncF plasmid replicon(s) present in the bacterial genomes. Of the collection of 72469 genomes, 5804 (8.01%) displayed discrepancies in the pMLST output.

Several situations may account for the discrepancies observed when multiple allele hits were detected. In one scenario, all of the alleles are present on one plasmid, thus the information about epidemiological background of a plasmid is not complete. The FAB formula created by the pMLST analysis of short-read sequences displays two different outputs varying between F31:A4:B1 and F36:A4:B1 as highlighted by the purple frame in Fig. 1. The detailed output may predict the presence of both of these alleles in all the versions of the tool together with a warning that multiple perfect hits were found (Fig. 1A). Regarding the long-read analysis, this situation also occurred for an allele variant combination F31/F36, either in a F31/F36:A4:B1, F31/F36:A-:B1 or F31/F36:A4:B58 plasmid. All the representative isolates carrying F31, F36 or both of these alleles (n=18) that underwent long-read sequencing confirmed carriage of these alleles on same plasmid. The allele combination F31/F36 was observed in 6.20% (360/5804) of all short-read sequenced isolates with multiple allele variants, however, frequency of carriage of these alleles on a single plasmid would have to be confirmed by long-read sequencing. The formula F31/F36:A4:B1 can be seen in isolate B0009108 with both of the FII allele variants being carried on a single plasmid (Fig. 1B). The FAB formula suggested by the pMLST tool differs between F31:A4:B1, F36:A4:B1 and F36:A4:B58 even though the correct output should state F31/F36:A4:B1/B58. The discrepancy between B1 and B58 allele variants seen in Fig. 1B is addressed later as the phenomenon occurs across various plasmids (Fig. 4).

**Figure 1.**
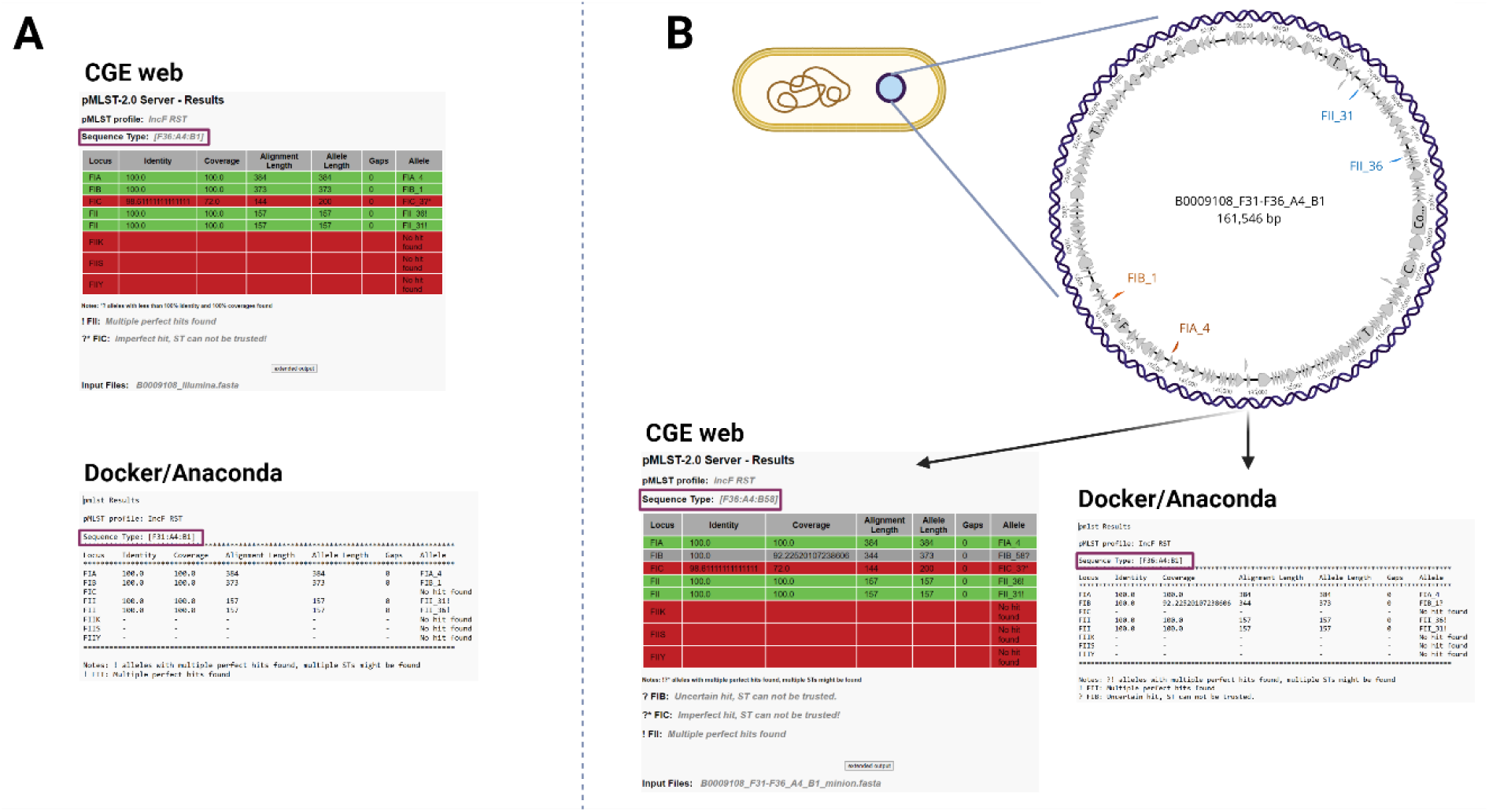
Presence of two FII alleles located on a single plasmid. Inconsistency between pMLST output of short-read sequences analysis using three versions of the tool with one of the FII allele variants being omitted while creating the FAB formula (A) and the results of long-read sequence analysis showing presence of an F31/F36:A4:B1 plasmid carrying both F31 and F36 alleles with an incorrectly assigned plasmid ST (B).

In the second situation, two or more alleles are located on different plasmids where it is unknown from short-read sequencing which plasmid is carrying specific alleles (Fig. 2). None of the analyses of short-read sequences pointed out the possibility of multiple plasmids occurring in the cell. This situation shows that the associations made between such plasmids and antibiotic resistance genes could be incorrect, since the true ST of the plasmid might be completely different than what is identified by *in silico* pMLST. This was observed in short-read sequences of isolate OV_CH_97 with pMLST output differing between F1:A2:B20 and F2:A2:B20 (Fig. 2A). However, the genomic analysis of long-read sequences of this isolate showed the presence of two IncF plasmids (Fig. 2B). After obtaining two different plasmids, pMLST was run on both of their sequences separately with two FAB formulas describing the respective plasmids as F1:A2:B20 and F2:A-:B-. When the pMLST was run on the complete long-read sequence only, the output did not differ from the one obtained by analysing short-read sequences with both of them leading to improper plasmid sequence type. A similar situation was observed in the isolate U23060 (Table S1) with both short and long-read analysis providing the FAB formula F29:A-:B10. However, after looking into the genome, two different plasmids F29:A-:B10 and K7:A-:B-were discovered with the FIIK allele being displayed only when running pMLST on the two plasmid sequences separately. Our analyses show that the pMLST tools are sometimes unable to identify the full plasmid cohort using both short-read, and long-read sequence data.

**Figure 2.**
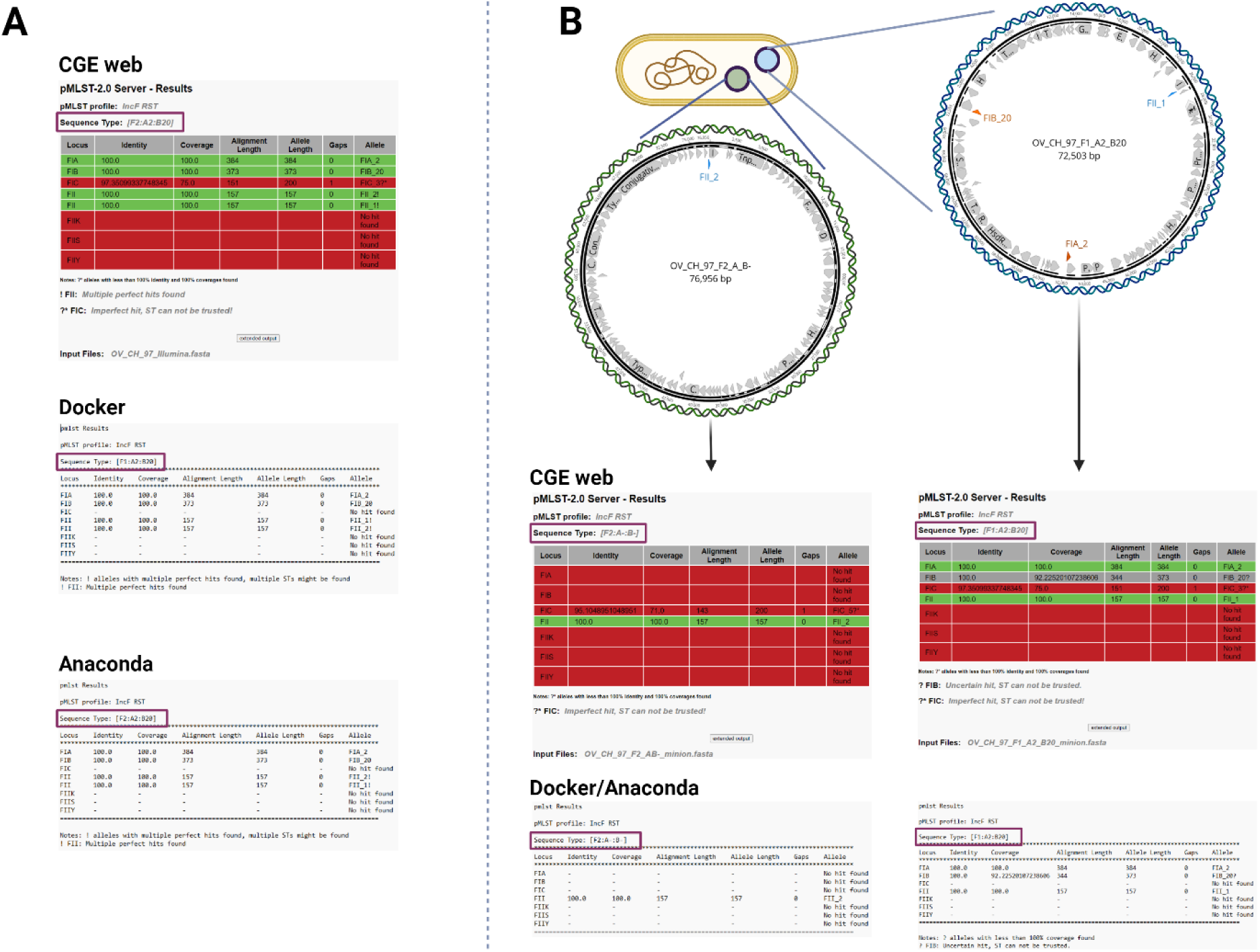
Presence of two IncF plasmids located in one cell with an incorrect FAB formula identification. The discrepancies observed during the analysis of the short-read sequences using three versions of pMLST which resulted in different plasmid STs for the same isolate (A), and the results obtained from long-read sequencing showing two plasmids, F1:A2:B20 and F2:A-:B-, being carried by the cell and a correct assignment of FAB formula (B).

In a third scenario, the start of the FII replicon overlaps with the end of the sequence of the FIC replicon, with both of them being located on a single plasmid (Fig. 3). Even though both of these alleles are present in the cell with 100% identity and 100% coverage, only one of them can take the position of FII allele in the FAB formula with the selection being random. The presence of both alleles was detected only by the CGE version of the pMLST tool, while the Docker and Anaconda versions omitted the FII allele. Short-read sequence of isolate B0009170 provided two pMLST outputs – FAB formulas C4:A-:B1 and F18:A-:B1 (Fig. 3A). Analysis of the long-read sequences showed both F18 and C4 allele variants present on a single C4/F18:A-:B1 plasmid with the alleles overlapping in one region (Fig. 3B). In this case, the C4 (200 bp sequence) and F18 (155 bp sequence) allele variants display an overlap of 43 bp, with the overlap shared among those alleles. Moreover, the allele variant F18 was detected only by the CGE web version in the analysis of both short-read and long-read sequences with Docker and Anaconda versions omitting this potentially crucial data. The same situation of overlapping alleles occurred in 5.48% (318/5804) of all short-read sequences carrying multiple allele variants and was connected to the C4/F18 allele combination.

**Figure 3.**
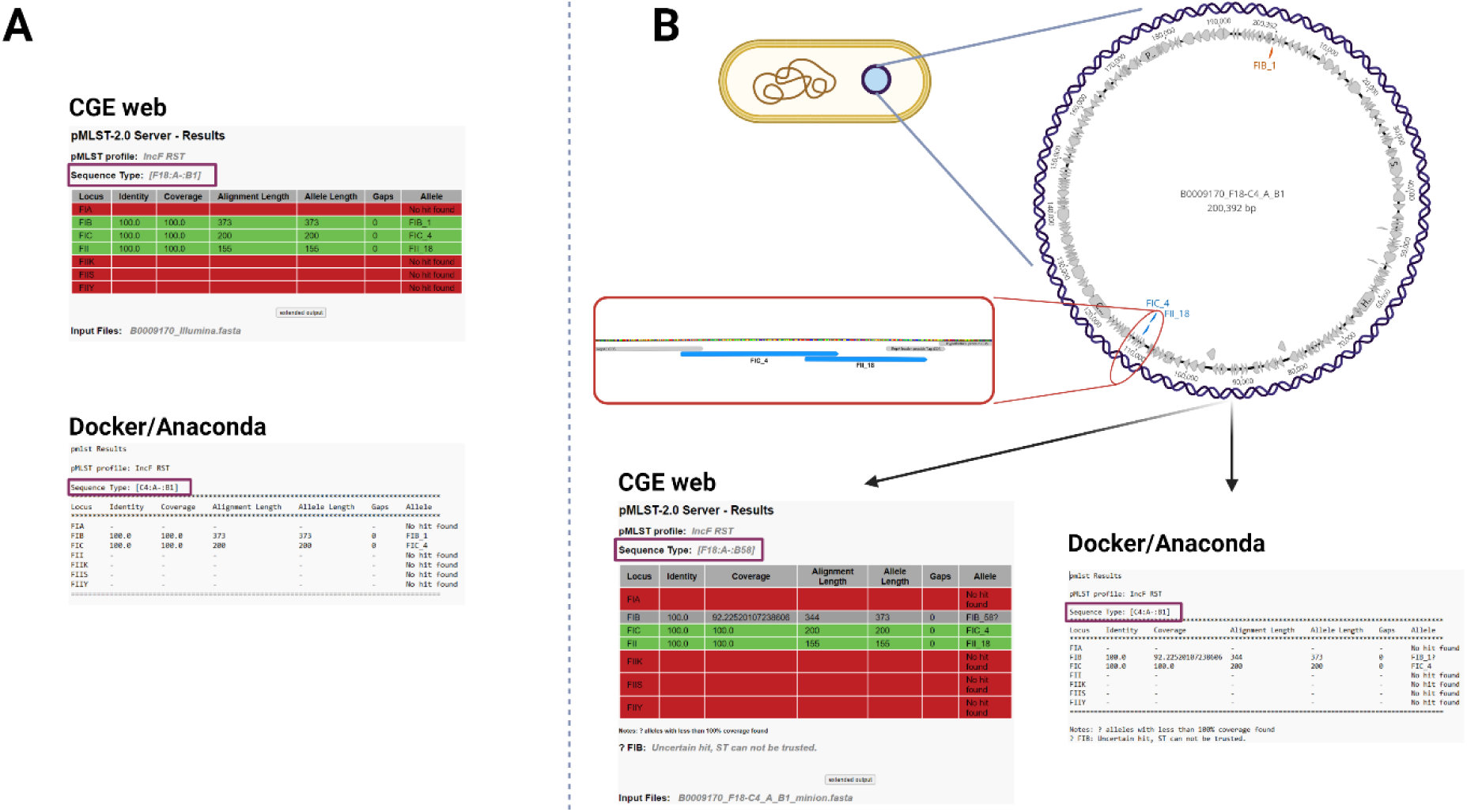
Presence of an FII and FIC allele overlapping one another on a single plasmid. Conflicting results obtained from short-read sequences using CGE web pMLST versions which showed the presence of the F18 allele while the Docker/Anaconda versions omitted it completely (A). Long-read sequences analysis displayed the same discrepancy between versions and showed a complete plasmid carrying both C4 and F18 allele with an overlap (B).

Lastly, discrepancies were found in the FIB allele variant of multiple RSTs (Fig. 4). This was observed in isolates B0009108 and B0009170 (see Fig. 1, Fig. 3) where the FAB formula showed either the allele B1 or B58. In short-read sequences, all of the pMLST tool versions correctly reported a B1 allele with 100% identity and 100% coverage but the pMLST analysis of complete plasmids resulted in randomly assigned B1 or B58 with 100% identity and 92.23% coverage. The alignment of these two FIB allele variants showed identical length and a single nucleotide difference located on the 11^th^ nucleotide of their sequence. In plasmids, the sequence of either one of these alleles starts 29 bp upstream of the replication protein. As there are only contigs and not circular genomes in *de novo* assemblies from short-read sequencing, the replication proteins along with the IncF allele sequences are usually in the middle of a contig. However, the long-read plasmid sequences are typically complete circular genomes, and thus are rotated during assembly to have the replication protein, i.e. the replication origin, at the beginning of a plasmid [12]. Since the pMLST tools work only with linear sequences, the circular plasmid sequence is cut at the replication origin, which leads to the 29 bp region at the beginning of the allele being omitted from the RST assessment resulting in the incorrect identification of a plasmid (Fig. 4).

**Figure 4.**
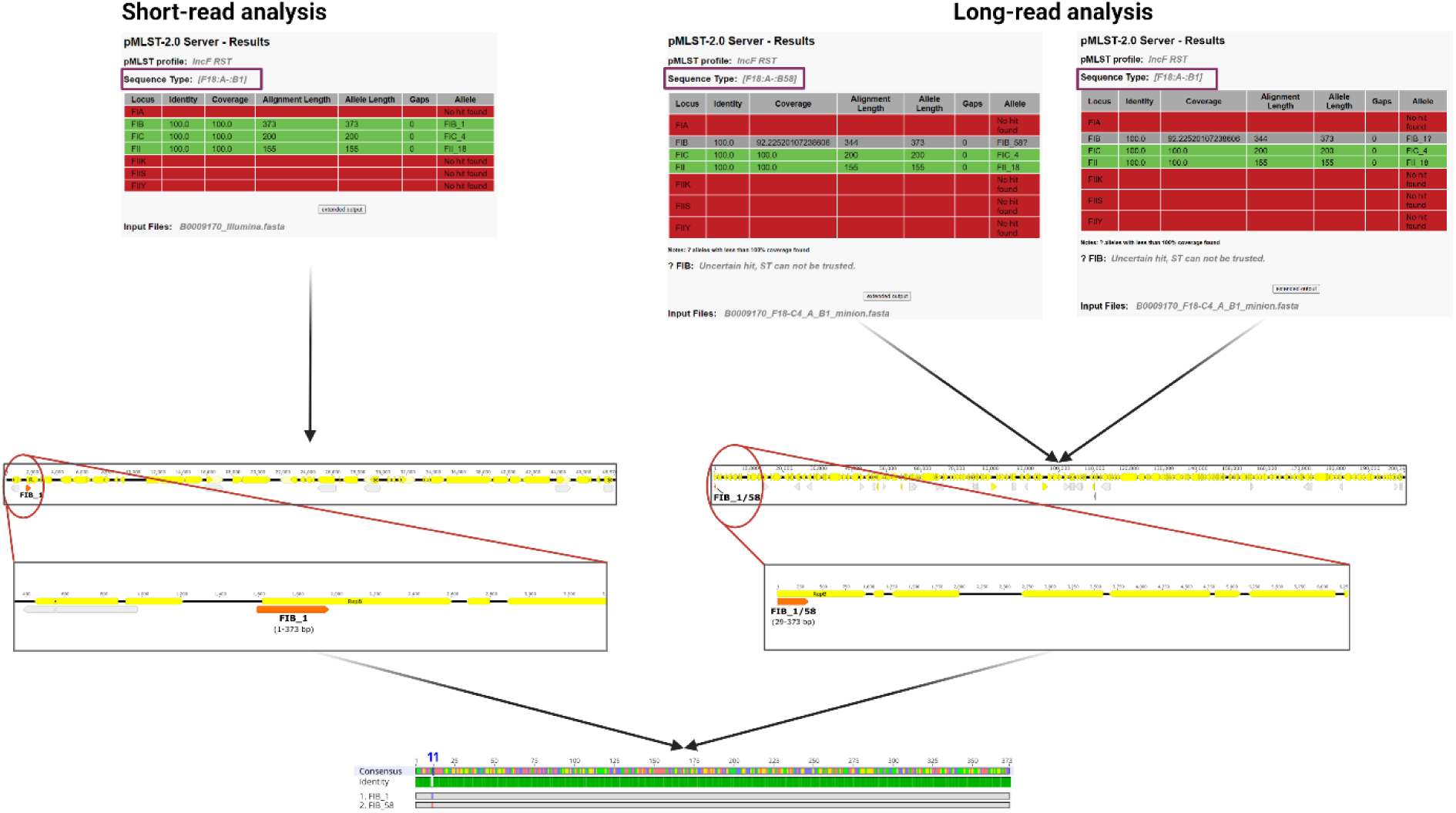
Discrepancies in the FIB allele variant observed between short-read and long-read analysis. Short-read analysis shows the presence of B1 allele with 100% coverage and 100% identity due to the complete allele being present in the middle of a contig. Repeated pMLST analysis of long-read assembly provided two different alleles – B1 and B58, which proved to differ in a single nucleotide located 11 nucleotides from the start of the allele sequence. Due to the circular nature of the complete plasmid, the sequence begins from the start of the replication protein causing the omission of the 29 bp region at the start of the allele, which contains the varying 11^th^ nucleotide of the allele sequence. All three pMLST tool versions provided the same results, therefore only the CGE web output is shown.

It is crucial to consider the detailed results of the pMLST tool and not only the automatically created FAB formula which does not contain the extended output to avoid potential omission of allele variants present in the genome. Such omission leads to incorrect pMLST results, likely biasing the evolutionary and epidemiological significance of specific plasmid lineages and their association with bacterial clones and ARGs.

### Most common combinations of the multiple allele variants

Out of 72469 bacterial genomes of *E. coli* of various STs as well as other Enterobacterales, 8.01% (n=5804) carried more than one allele variant of an IncF replicon (Table S1). Even though the phenomenon of multiple allele variants was linked mainly to the FII replicon (99.00%; 5746/5804), it was sporadically observed also in the FIA (0.90%; 52/5804) and the FIB replicons (0.40%; 23/5804). Small number of genomes (0.29%; 17/5804) carried multiple allele variants in more than one allele simultaneously, usually in the FII allele and then either in the FIA or FIB. Number of various allele variant combinations reached up to 900. The most common combinations observed in the short-read sequences of the whole collection were F31/F36 (6.20%; 360/5804), C4/F18 (5.48%; 318/5804), F1/F2 (4.72%; 274/5804) and F2/F29 (3.84%; 223/5804) (Fig. 5). Mistaking some of these alleles for their counterparts might obscure epidemiologically important insights. This was notable for the C4/F18 and F2/F29 allele combinations. The F18 allele often occurs in plasmids of the ColV group carrying large number of virulence genes and associated with avian pathogenic *E. coli* (APEC) clones in poultry [13], [14], [15]. The F29 allele constitutes one part of the F29:A-:B10 (also known as pUTI89-like plasmid) which is known for its conservative virulence region *cjrABC*-*senB*. This plasmid is commonly linked to *E. coli* strains causing urinary tract infections, often with nosocomial potential [16], [17]. Improper pMLST reporting might lead to omission of important epidemiological insights regarding these and other strains and associated virulence and/or resistance plasmids. As such caution is advised when using these tools alone to infer host strain phylogeny.

**Figure 5.**
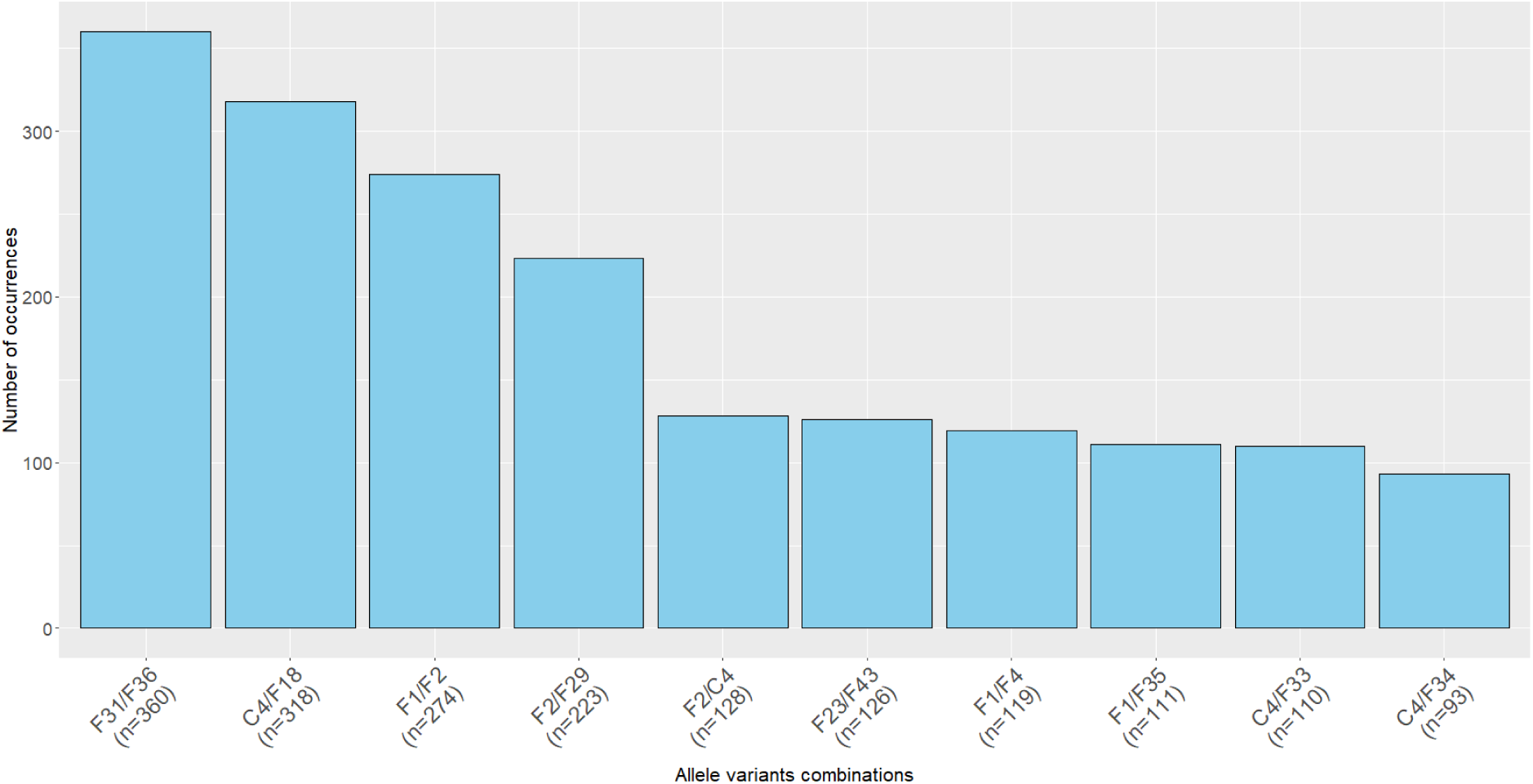
The most common allele variants combinations obtained by short-read sequencing of the complete collection of 5804 analysed Enterobacterales genomes carrying multiple allele variants

This phenomenon has not been described before except for a few papers where the multiple allele variants appear in the output but are not commented on [18], [19]. It was previously reported in various *E. coli* STs such as ST10, ST58, ST155, ST963, ST973 and ST1140 as well as in the ST we focused on the most, ST131 [18]. The improper allele assignment in these cases regarded only the FII allele, which was also the most common incorrectly assigned allele in our collections. Even though various combinations were present, the F29 allele variant was the most prevalent one occurring in such cases which highlights the importance of proper assignment due to its association with pUTI89-like plasmid. Combinations of other variants were observed in a lower number of cases, none of them corresponding to the most prevalent combinations in our results [18], [19].

Our findings were supported by analysing the real prevalence of multiple allele variants carried on a single plasmid in the collection of 7441 complete sequences of IncF plasmids obtained from GenBank. In this dataset, the prevalence was distinctly lower (3.52%, 262/7441), which matches the long-read sequencing results obtained in this study, where we saw that even though the FAB formula from short-read analysis looks like all the replicons are located on one plasmid, it is not always the case (Fig. 2).

## Conclusion

The study identified discrepancies between results reported by all versions of *in silico* pMLST tools and the real biological situation in the bacteria. Even though the tool provided us with all the IncF allele variants present in the cell, there was no possibility of assigning them to their respective plasmids. FAB formulae were also created by a random selection of these IncF alleles excluding the presence of their multiple variants. Since the STs of plasmids are often linked to specific bacterial lineages, improper or insufficient pMLST report might lead to the omission of certain epidemiologically relevant plasmid-bacterium-antibiotic resistance associations.

## Acknowledgements

We would like to thank Ivana Jamborova, Martina Masarikova, Alois Cizek, Aneta Papouskova and Patrik Mlynarcik for providing their sequencing data together with metadata.

## Authors’ contributions

MR, JP, IP and MLC analysed the sequencing data. JP and MLC obtained sequence data from public databases. MR performed the long-read sequencing. SPD and MD provided valuable insights and supervised the project. MR wrote the manuscript and all other authors contributed to revisions. The authors read and approved the final manuscript.

## Transparency declarations

None to declare.

